# Overcoming host restrictions to enable continuous passaging of human noroviruses in human intestinal enteroids

**DOI:** 10.1101/2025.05.22.655571

**Authors:** Gurpreet Kaur, Sue Crawford, Sasirekha Ramani, BV Ayyar, Aaya Boussattach, Xi-Lei Zeng, Robert L. Atmar, Hoa Nguyen Phuc, Cristian Coarfa, Mary K. Estes

**Author notes:** Address correspondence to Mary K. Estes.

## Abstract

Human noroviruses (HuNoVs), the leading cause of viral gastroenteritis, can now be cultivated in human intestinal enteroids (HIEs). However, indefinite passaging of HuNoVs in HIEs remained a challenge, necessitating the use of patient stool samples as viral inocula. Using RNA-seq, we identified host restriction factors that might limit viral passaging. CXCL10, CXCL11, and CCL5 were among the most upregulated chemokines, suggesting their potential as host restriction factors. TAK-779, a CXCR3/CCR5/CCR2 antagonist, enhanced GII.3 HuNoV replication and viral spread in a dose- and time-dependent manner, enabling successful passaging of GII.3 HuNoV in two different HIE lines and generation of viral stocks. TAK-779 also enhanced replication of GI.1 and GII.17 strains, but not GII.4, suggesting strain-specific host interactions or immune evasion. This breakthrough in passaging provides critical insight into HuNoV-host interactions, establishes a scalable in vitro system for virus propagation, and opens avenues for structural, biochemical and therapeutic studies.

## Introduction

Human noroviruses (HuNoVs), members of the *Caliciviridae* family, are single-stranded, positive-sense RNA viruses and a leading cause of acute viral gastroenteritis worldwide. HuNoV infection is characterized by vomiting and diarrhea, with severe outcomes more prevalent among young children, the immunocompromised, and the elderly (*1–4*). In individuals with compromised immune systems, infections can become chronic (*5–7*). Annually, HuNoVs are responsible for approximately 677 million cases of diarrhea and over 213,000 deaths globally (*8*). In the United States alone, these infections result in an estimated 109,000 hospitalizations, imposing an economic burden exceeding $10.6 billion in healthcare and societal costs (*9, 10*). Despite the substantial public health impact, there are currently no FDA-approved vaccines or antiviral therapies for HuNoV with current management strategies relying solely on supportive care, including fluid and electrolyte replacement (*11, 12*).

Although the first documented outbreak of HuNoV was reported in 1968 (*13*), a major limitation for HuNoV research for nearly five decades was the absence of a robust, reproducible cultivation system. Several distinct *ex vivo* HuNoV culture models including BJAB cells (*14, 15*), zebrafish (*16, 17*), human induced pluripotent stem cell-derived (iPSC) intestinal organoids (*18*), transformed salivary cell lines (*19*), and tissue stem cell-derived human intestinal enteroids (HIEs) (*20*) have been reported in the last decade. Among these, HIEs closely recapitulate the intestinal tropism observed in infected persons (*21*) and support reproducible replication of multiple HuNoV strains with high-yield (*22, 23*). Furthermore, HIEs from different donors effectively model epidemiological patterns and confirm the genetic susceptibility associated with HuNoV infection (*20, 24*).

Since the first publication in 2016, the HIE system has been adopted by multiple laboratories worldwide for diverse HuNoV studies and have supported investigations into mechanisms that regulate viral infection (*24–29*), testing antiviral therapies (*30–35*), viral inactivation (*36–40*), antibody neutralization potential (34262046 *41, 42–48*), and viral persistence in the environment (*49, 50*).

One major obstacle that persisted in HIEs was the lack of indefinite serial passaging of virus. Our previous studies demonstrated that HIEs support up to four consecutive passages of GII.4 HuNoV with low and decreasing yields for each passage (*20*). Currently, no in vitro culture system allows for indefinite HuNoV propagation, requiring researchers to rely on clinical stool specimens for infection studies. However, the limited volume of clinical specimens and variability in virus titers between stool samples pose challenges for research that is comparable across laboratories. The lack of sustainable laboratory stocks also limits the establishment of high-throughput screening platforms involving thousands of compounds for antiviral drug discovery. Currently, antiviral testing is feasible but requires significant effort, even for small-scale compound libraries (∼300 compounds), due to the limited availability of HuNoV-positive stool samples (*12*).

Thus, establishing an efficient HuNoV culture system that allows serial passaging, and the propagation of viral stocks is important. We evaluated the role for uncharacterized host factors that limit or promote viral replication to address challenges with indefinite passaging. In previous investigations into host mechanisms restricting virus replication, we and others identified a key role for interferons (IFNs) and the JAK-STAT signaling pathway. We found replication of the GII.3 HuNoV strain was significantly enhanced in STAT1-knockout HIEs, which exhibited increased permissiveness and viral spread (*29*). This finding was supported by another study demonstrating enhanced HuNoV replication through targeted inhibition of Janus kinase 1 (JAK1) and Janus kinase 2 (JAK2) (*27*). Additionally, genetic engineering of secretor-negative, non-susceptible HIEs to render them secretor-positive and susceptible to infection resulted in enhanced viral replication of multiple strains (*22, 24*). Further optimization of the culture system, including the use of commercially available media, has also led to increased viral replication of multiple strains (*22, 23*). Despite these advancements, the challenge of sustained passaging remained unresolved. In this study, we characterized host factors that continue to restrict viral replication in genetically engineered HIEs that exhibit enhanced permissiveness. By comparing these HIEs to their parental counterparts, we identified key determinants limiting sustained HuNoV propagation and developed a strategy for optimizing long-term viral cultivation of GII.3 HuNoV.

## Material and methods

### Maintenance and culture of HIEs

HIE cultures used in this study were obtained from an HIE bank maintained by the Gastrointestinal Experimental Model Systems (GEMS) Core of the Texas Medical Center Digestive Diseases Center (TMC DDC). Establishing HIE cultures from human surgical or biopsy tissue was approved by the Baylor College of Medicine (BCM) Institutional Review Board as described previously (*20, 23*). J2 (*20*) and the genetically-modified J4FUT2-KI (*24*) and J8FUT2-KI in which the *Fucosyltransferase 2* gene was knocked into the parental J4 and J8 HIEs using CRISPR-Cas9 and the and the J2STAT1-KO HIE, in which STAT1 was knocked out of the J2 HIE using CRISPR-Cas9 guide RNAs (*29, 51*) were grown as multilobular 3-dimensional (3D) HIEs in Matrigel and maintained in L-WRN complete media as described (*23*). For HuNoV infection, the 3D HIEs were dissociated by trypsinization and pipetting and seeded onto collagen IV coated 96-well plates using a 1:1 ratio of IntestiCult™ Organoid Growth Medium Human Basal Medium (StemCell Technologies, 100-0190) and Organoid Supplement (StemCell Technologies, 100-0191) (OGM proliferation medium) supplemented with 10 µM ROCK inhibitor Y-27632 for 24 hours. The HIEs were then differentiated for 5 days using a 1:1 ratio of Intesticult™ OGM and complete medium without growth factors (CMGF-medium, advanced DMEM/F12 prepared with 1X GlutaMAX and 10 mM HEPES) (OGM differentiation medium) with media changes every two days prior to inoculation with HuNoV (*22, 23, 52, 53*).

### Virus filtrates and HuNoV infection

Ten percent stool filtrates containing HuNoV were prepared as described previously (*20, 23*). Briefly, 4.5 ml of ice-cold phosphate-buffered saline (PBS) was added to 0.5 ml of stool, homogenized by vortexing, and sonicated three times for 1 min. The sonicated suspension was centrifuged at 1,500 × g for 10 min at 4°C. the supernatant was transferred to a new tube and centrifuged a second time. The resulting supernatant was passed serially through 5-μm, 1.2-μm, 0.8-μm, 0.45-μm, and 0.22-μm filters depending on stool texture. The filtered sample was frozen in aliquots at −80°C until use. Virus stool filtrates are described in Table 1.

**Table 1.**
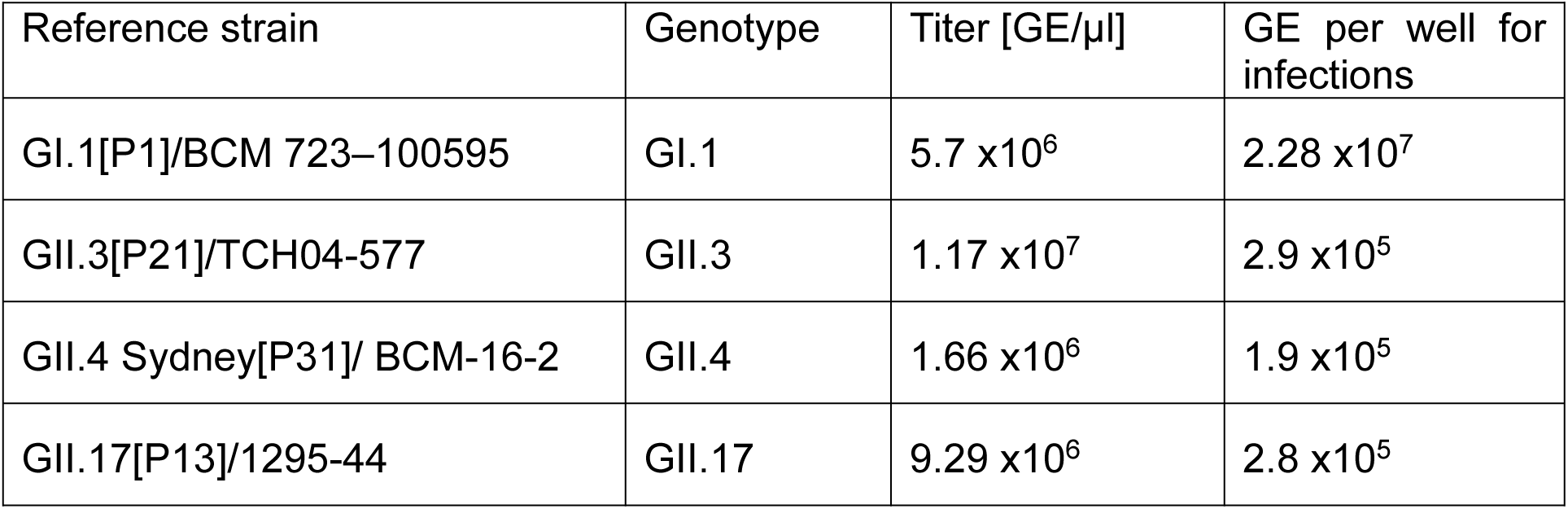
List of HuNoV isolates used in this study and their infectivity to HIE lines.

Five-day differentiated HIE monolayers were washed once with CMGF (−) medium (*22, 23*) and inoculated with HuNoV for 1–2 h at 37°C. The inoculum was removed, and monolayers were washed twice with CMGF (−) medium to remove unbound virus. OGMd differentiation medium (100 μL containing 500 μM bile acid glycochenodeoxycholic acid [GCDCA]) was then added, and the cultures were incubated at 37°C for the indicated time points.

### RNA-seq analysis

We performed RNA-seq analysis on three 5-day differentiated jejunal HIE lines, parental J2 and the genetically-modified J2STAT1-KO and J4FUT2-KI, inoculated with gamma-irradiated GII.3 (gGII.3) or with infectious GII.3 at high MOI (3.2 × 10^7^ genome equivalents/well). Total cellular RNAs were extracted from GII.3- or gGII.3-inoculated monolayers at 10, and 24 hours post infection (hpi) using the RNeasy Mini Kit according to the manufacturer’s protocol (Qiagen). The RNA integrity was assessed on a 2100 Bioanalyzer (Agilent). RNA sequencing library preparation was performed at BCM following a standard protocol as previously described (*29*). Libraries were created from the cDNA by first blunt ending the fragments, attaching an adenosine to the 3′ end, and finally ligating unique adapters to the ends for PCR amplification. Paired end sequencing (2 × 150 bp) was performed on an Illumina HiSeq 2500 for a depth of approximately 43 million paired end reads per RNA sample. Raw sequence reads were checked for quality using the FASTQC package ver. 0.11.9 and reads were trimmed with TrimGalore ver. 0.6.5 with default settings for adaptive trimming and for base quality filtering. To confirm infection with GII.3, a custom reference comprised of the human genome build hg38 and the GII.3 genome was constructed using STAR (*54*), then the trimmed reads were mapped to this new reference using STAR, and gene expression was quantified using featureCounts (*55*). Gene expression for all genes was normalized to counts per million (CPMs), then gene expression for GII.3 HuNoV genes ORF1, ORF2, and ORF3 was plotted using GraphPad Prism. For further gene expression analysis, trimmed reads were aligned to human genome build GRCh38.98 using HiSAT2 ver 2.2.1 (*56*) and a count matrix was generated from the aligned reads using featureCounts. Enriched pathways were determined using Gene Set Enrichment Analysis (GSEA) (*57*).

### TAK-779 treatment

TAK-779 was purchased from Medchem Express (Cat. No. HY-13406) and reconstituted in water to make 5 mM stocks and frozen in single use aliquots to avoid repeated freeze-thaws. HIE monolayers were pre-treated with different doses of TAK-779 or vehicle control in OGM differentiation media for 3 h. Monolayers were inoculated in media with vehicle control (water) or TAK-779 and with stool filtrate containing genome equivalents (GEs) of HuNoV per well as indicated in Table 1 for 1 to 2h (binding time increased to 2 h in GI.1 HuNoV experiments) at 37°C in 5% CO2. Monolayers were then washed twice with CMGF (-) to remove unattached virus. The inoculated cells were maintained in OGM differentiation media with vehicle control or TAK-779 for indicated time points.

### Tissue culture infectious dose 50% (TCID_50_) assay

We determined GE/TCID50 values to allow us to evaluate the permissiveness of the J4FUT2-KI HIE line for infection with GII.3 virus in the presence of TAK-779 compared to vehicle control. We determined the TCID50 per inoculum by examining the numbers of wells that showed increases in GE of virus per well at 48 h above baseline (at 1 h) as measured by RT-qPCR (*22, 30, 58*). TCID50 per volume was calculated by Reed-Muench. We also used RT-qPCR to measure the number of GEs per volume using a genogroup-specific standard. We then calculated the GE per TCID50. Compiled data from three independent experiments are presented. In this context, the information is expressed as the number of GEs (log10[GE]) per TCID50 at which 50% of the cultures are infected as determined by RT-qPCR.

### Quantification of viral replication by RT-qPCR

RNA extraction and RT-qPCR were performed as previously described (*23*). In brief, total RNA was extracted from each infected well using the KingFisher Flex purification system and MagMAX-96 Viral RNA isolation kit. RNA extracted at 1 hpi was used as a baseline to determine the amount of input virus that remained associated with cells after washing the infected cultures to remove unbound virus. The primer pair and probe COG2R/QNIF2d/QNIFS (*59*) were used in the RT-qPCR for detection of GII genotypes, and the primer pair and probe NIFG1F/V1LCR/NIFG1P (*60*) were used for GI.1. RT-qPCR was performed with qScript XLT One-Step RT-qPCR ToughMix reagent with ROX reference dye (Quanta Biosciences) in an Applied Biosystems StepOnePlus thermocycler. Each extracted RNA was run in duplicate with the following cycling conditions: 50°C (15 min), 95°C (5 min), followed by 40 cycles of 95°C (15 s) and 60°C (35 s). Standard curves based on recombinant GII HuNoV RNA transcripts were used to quantitate viral GEs in RNA samples. The limit of detection of the RT-qPCR assay was 20 GEs. Samples with RNA levels below the limit of detection of the RT-qPCR assay were assigned a value that was one half the limit of detection (10) of the assay. A threshold for successful viral replication was established by considering a 0.5 increase in log10(GE) after 24 hpi relative to the genomic RNA detected at 1 hpi (*23*)

### Immunofluorescent detection of HuNoV-infected HIEs

Infected monolayers in 96-well plates were washed with 1X PBS and fixed with 100% methanol at -20°C for 20 min, washed with 1X PBS three times, blocked with 5% BSA in 1X PBS at room-temp for 1 h and incubated with primary antibodies (1:1000 dilution of guinea pig anti-HuNoV polyclonal Ab, 1:150 dilution of rabbit anti-RdRp, 1:200 dilution of mouse anti-VPg) in 3% BSA in 1X PBS at 4°C overnight. The monolayers were then washed with 1X PBS four times incubated with 1:500 dilution of goat anti-rabbit 488 donkey anti-mouse 549 (#610-742-124, Rockland), anti-rabbit 649 (#611-743-127, Rockland), and anti-guinea pig 488 secondary antibodies (#606-141-129, Rockland) for 2 hrs at 4 °C. The cells were washed three times, and nuclei were stained with 4, 6-diamidino-2-phenylindole (DAPI) (300 nM) before visualizing the viral proteins. Z-stack images were captured using a Zeiss Laser Scanning Microscope LSM 980.

### Passaging of HuNoVs

J4FUT2-KI and J8FUT2-KI HIEs were pretreated with 30 µM TAK-779 or vehicle control for 3 hours at 37°C in 5% CO₂. Following pretreatment, monolayers were inoculated in OGM differentiation media containing 500 uM GCDCA supplemented with either vehicle control or 30 µM TAK-779, along with GII.3 HuNoV stool filtrate containing approximately

2.9 × 10⁵ GEs of HuNoV per well. The inoculation was carried out for 1 hour at 37°C in 5% CO₂. Post-infection, monolayers were washed twice with CMGF (-) medium to remove unattached virus and subsequently maintained in OGM differentiation media with 500 µM GCDCA supplemented with either vehicle control or 30 µM TAK-779 for 96 hours. Cells and supernatants were harvested and subjected to sonication (3 cycles of 1 min on and 1 min off on ice). Lysates were then centrifuged at 400 × g for 10 minutes to remove cellular debris. The resulting supernatant and pellet fractions were quantified using heat release method (*61*). Samples were serially diluted (1:5, 1:10, 1:50, and 1:100), ensuring a final volume of 50 µL and then heated at 95°C for 5 minutes and immediately transferred to ice. GEs were quantified using a standard RT-qPCR protocol as described above. Monolayers were then inoculated using supernatants containing 1–5 × 10⁵ GE of passaged GII.3 HuNoV per well, following the same infection protocol. It was crucial to accurately titer the input virus and use a low multiplicity of infection (MOI) to avoid triggering an early immune response. After 1 hour of incubation at 37°C in 5% CO₂, monolayers were washed twice with CMGF (-) and maintained in OGM differentiation media containing 500 µM GCDCA with either vehicle control or 30 µM TAK-779 for 96 hours.

### Statistical analysis

Each experiment was performed two or more times, with three technical replicates of each culture condition and time point. Two technical replicates were performed on each well for RT-qPCR and then averaged. The GEs between the three technical replicate wells per condition were then averaged. Data from each averaged condition were then pooled among repeated experiments. The number of experiments (n) is noted in the figure legends. All statistical analyses were performed on GraphPad Prism for 381 Windows (GraphPad Software, La Jolla, CA, USA). Comparison between groups was done using a two-way analysis of variance (ANOVA) and Sidak post hoc multiple comparisons analysis. TCID50 comparisons were done using Student’s *t*-test, *P* values of <0.05 were considered statistically significant.

## Results

### RNA sequencing reveals epithelial responses to GII.3 infection differ between HIEs, with cytokine signaling in immune system as a common response

To evaluate intestinal epithelial responses to HuNoV that might restrict replication, we infected monolayers of three susceptible, jejunal HIE lines: J2 and two genetically-modified cultures, J2STAT1-KO and J4FUT2-KI, with the bile acid-dependent GII.3 virus (Fig. 1A). Gamma-irradiated (gGII.3) virus was used as a control to determine any effect of stool alone on HIEs. We have previously shown that knocking out STAT1, a key transcription factor regulating IFN signaling, enhances GII.3 HuNoV replication accompanied with viral spreading (*29*). The J4FUT2-KI HIE line was generated by genetically knocking in the *fucosyltransferase 2* (*FUT2*) gene into the J4 HIE line, which is genetically and phenotypically secretor negative and not susceptible to infection by many HuNoV strains (*24*). In addition, we have shown that the J4FUT2-KI HIE line supports better replication of multiple HuNoVs (*22*). The parental J4 HIEs do not support GII.3 replication at 24 h hpi and were not used in the RNA-seq analysis. Transcriptional responses to GII.3 inoculation were analyzed at 24 hpi because we previously reported no detection of differentially regulated genes at early time points (6 and 10 hpi) (*29*). The cultures were infected based on evaluating replication by increased viral RNA at 24 hpi compared to 10 h binding (Fig. S1A) and enhanced reads of the expressed viral genes in the RNA-seq dataset (Fig. S1B). We previously demonstrated that the genetic background of the HIE cultures impacts analyses of transcriptional signatures to viral infections, with responses first clustering by HIE line (*29*). Consistent with this, principal component analysis (PCA) showed that HIE line was the largest driver of variance (Fig. 1B).

**Figure 1.**
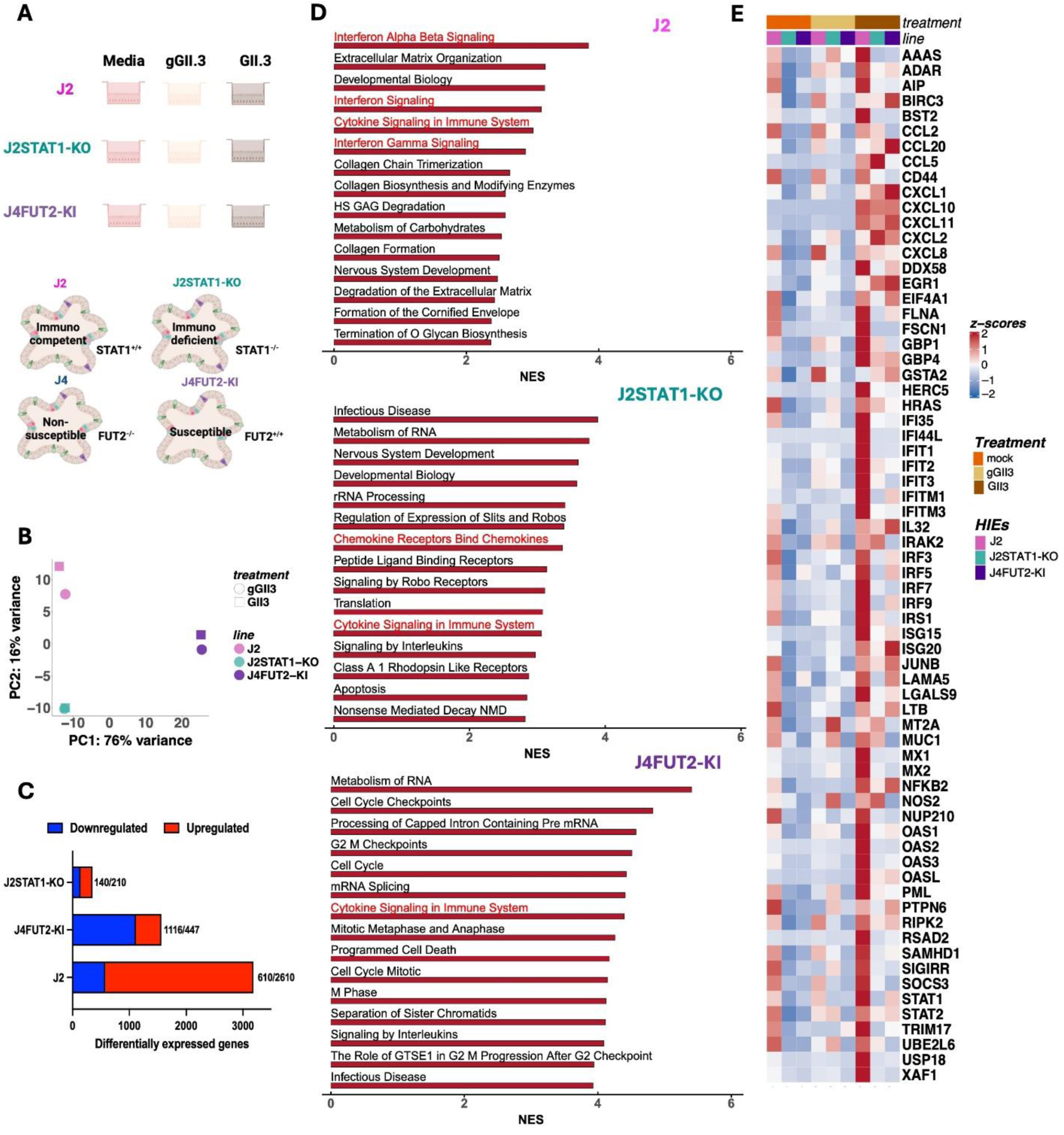
Epithelial responses to Gil.3 infection differ between HIEs with cytokine signaling in immune system as a common response. (A) Schematic of infection for RNA-seq analysis. Monolayers (J2, J2STAT1-KO and J4FUT2-Kl) of HIEs were inoculated with gamma-irradiated Gil.3 (gGII.3), media or infectious Gil.3 stool filtrate. Cultures were harvested at 24 hpi. (B) Principal component analysis (PCA) was performed on normalized log2 counts per million (cpm). (C) The increased or decreased genes (Gll.3-infected over gGII.3-inoculated cultures at 24 hpi, fold change of at least 1.5x) are summarized in the bar chart. (D) Gene set enrichment analysis (GSEA) results using the Reactome Biological Processes compendium showing the top 15 common upregulated pathways for the Gil. 3-infected over gGII.3-inoculated cultures at 24 hpi in the three different HIEs, with immune pathways indicated in red. (E) ISGs selected from the core genes contributing to the upregulated pathways shown in D (in red) are visualized in a heatmap.

Comparison of the level of differentially expressed host genes in GII.3-infected cultures to gGII.3-treated cultures showed, in concordance to the PCA plot, GII.3 HuNoV infection induced changes in transcriptional responses in the three lines. However, the number of genes that were altered between GII.3-infected and gGII.3-treated HIEs varied between the HIE lines with the highest number of different genes seen in the J2 followed by the J4FUT2-KI and then the J2STAT1-KO HIEs (Fig. 1C). These findings showed, for the first time, a reduced host transcriptional response to GII.3 infection in genetically modified lines compared to the unmodified J2 line.

Next, we used gene set enrichment analysis (GSEA) to determine the key pathways involved in the response to GII.3 infection in the different HIE lines. Controlling for the effect of potential immune modulators in stool using gGII.3, our GSEA analysis revealed that the top 15 pathways up regulated by GII.3 infection were primarily IFN signaling and antiviral responses linked to in interferon signaling genes (ISGs) in the parental J2 line as previously reported for GII.4 HuNoV infection of HIEs (Fig. 1D). Unexpectedly, in the modified J2STAT1-KO and J4FUT2-KI lines, cytokine signaling in the immune system and chemokine-mediated antiviral responses emerged as the dominant immune-related upregulated pathways. Additionally, pathways related to cell cycle regulation and translation were also enriched in the genetically-modified lines, suggesting a distinct host response in these lines compared to J2. Consistent with these findings, we observed a robust increase in expression of ISGs as shown in a heat map (Fig. 1E) that included genes such as IFIT1, IRF3, BST2, RSAD2, CXCL10, CCL5, CXCL11, HERC5, IFITM1, ISG15, STAT1 in GII.3-infected J2 HIEs at 24 hpi, as well as other antiviral genes OASL and USP18. By contrast, CXCL10, CXCL11, CCL5, CX3CL1 are increased in the J2STAT1-KO HIEs. In addition, CXCL10, CXCL11, CXCL1, CXCL2, and CCL20 were upregulated in J4FUT2-KI HIEs. In summary, the transcriptome data indicate that GII.3 HuNoV infection of the J2 HIE line induces a robust IFN-mediated ISG response. In contrast, GII.3 infection of the genetically-modified J2STAT1-KO and J4FUT2-KI lines induces a chemokine response, which lead us to hypothesize that inhibiting these host factors may allow us to passage GII.3 in the genetically-modified HIE lines.

### TAK-779 Enhances GII.3 HuNoV Replication in a Dose-Dependent Manner in HIE Lines

We hypothesized that the detected chemokine signaling in the genetically-modified cultures might represent a primary immune barrier restricting HuNoV replication. To investigate the role of chemokine signaling in restricting HuNoV replication, we evaluated the effect of TAK-779, a CCR5/CXCR3/CCR2 antagonist (*62*), on viral replication in three HIE lines. HIEs were pre-treated with TAK-779 for 3-hours before infection to account for potential early immune activation by stool-derived components. The compound was also added during and following infection. Cytotoxicity assays (LDH assay) determined that TAK-779 is cytotoxic at concentrations exceeding 30 µM in J2 and J4FUT2-KI HIEs, while J2STAT1-KO cells exhibited toxicity even at lower concentrations and were not used in further experiments (Fig. S2). We found TAK-779 enhances GII.3 HuNoV replication in a dose-dependent manner, with the highest enhancement [an approximately 1-log10 increase in genome equivalents (GE)] observed at 30 µM in J2 and J4FUT2-KI cells at 48 hpi (Fig 2A and 2B).

**Figure 2.**
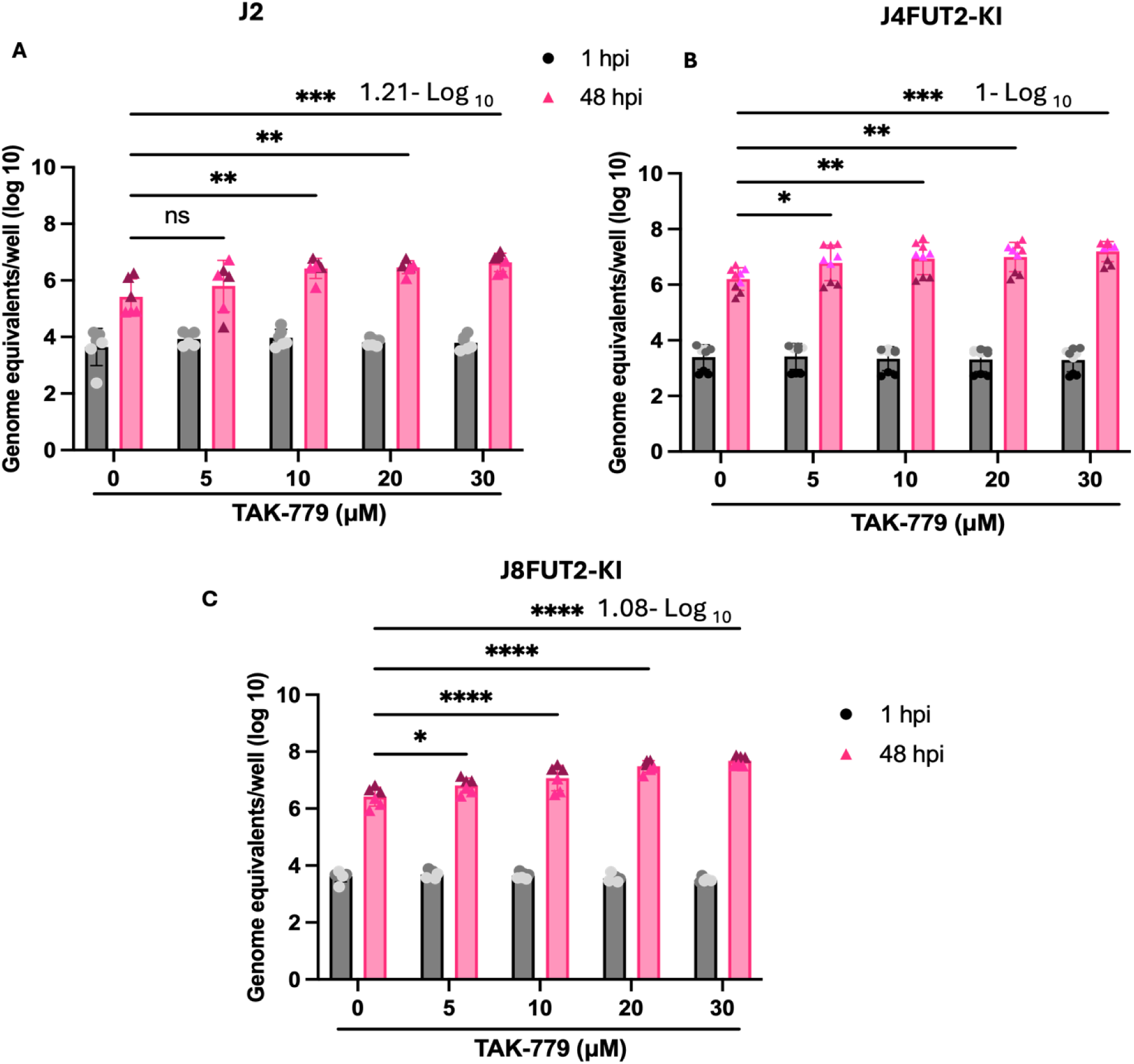
The CCR5/CXCR3/CCR2 antagonist TAK-779 enhances GII.3 replication in a dose-dependent manner in three HIE lines. HIEs were pre-treated for 3 hrs with the indicated concentrations of TAK-779 and then infected with GII.3 HuNoV (2.9 x 10° GEs/well) in the absence or presence of the same concentrations of TAK-779. After washing, the cells were cultured for 48 hrs at 37°C in the absence or presence of TAK-779. Viral GEs at 1 and 48 hpi were quantified by RT-qPCR. Mean data compiled from two independent experiments (J2 and J8FUT2-KI HIEs) and three independent experiments (JAFUT2-KI HIEs) with three wells per experiment are shown; error bars show SD. Experiments are denoted with different symbol colors. Significance was determined using one-way ANOVA comparing 24 h replication, no TAK-779 to each concentration of TAK-779 (* p value < 0.05; ** p <0.01; *** 0 <0.001; ****,0.0001).

To further validate this effect, we tested an additional HIE line, J8FUT2-KI, which was engineered based on the same strategy use for the J4FUT2-KI. Viral replication was enhanced in a similar manner in the J8FUT2-KI, with TAK-779 exhibiting the most pronounced effect at 30 µM (Fig. 2C). Notably, the one log10 increases in replication in the presence of TAK-779, compared to no inhibitor, represents the highest level of replication achieved in our HIE culture system for GII.3 HuNoV.

### Pre- and Post-Treatment with TAK-779 Maximizes GII.3 HuNoV Replication

We next sought to investigate whether the observed enhancement results principally from priming the host cells with TAK-779 or whether extending the pre-treatment duration could further augment viral replication. To determine the optimal conditions for TAK-779-mediated enhancement of GII.3 HuNoV replication, multiple treatment regimens were evaluated: (a) pre-infection treatment with 30 µM TAK-779 for 3 hours without post-infection treatment, (b) post-infection treatment with TAK-779 for 48 hours, (c) combined pre-infection treatment (3 hours) and post-infection treatment (48 hours), and (d) extended pre-infection treatment (24 hours) followed by treatment for 48-hour post-infection. While all treatment conditions significantly enhanced GII.3 replication compared to no treatment alone, the most robust enhancement was observed with the combined pre- and post-infection treatment condition, indicating a combined, synergistic effect of TAK-779 in promoting viral replication (Fig 3). This was seen in a dose-response analysis (Fig. S3). Notably, extending the pre-infection treatment duration to 24 hours did not result in further enhancement, suggesting that a 3-hour pre-infection treatment followed by 48-hour post-infection treatment is the optimal tested condition that produced the maximal (1.5 log10 increase) GII.3 HuNoV replication (Fig 3).

**Figure 3.**
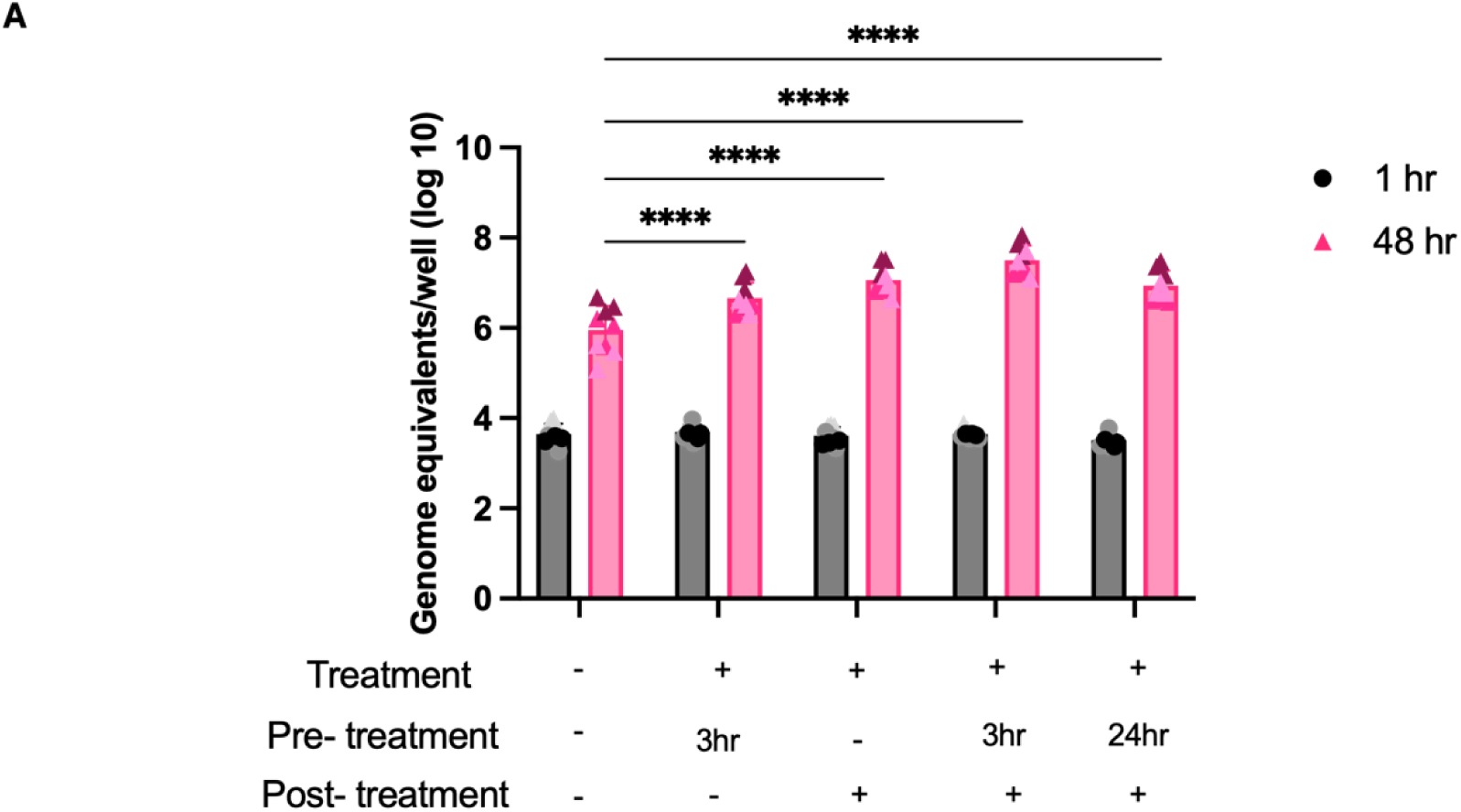
Pre- and post-treatment with the CCR5/CXCR3/CCR2 antagonist TAK-779 results in the best enhancement in GII.3 replication. Different conditions, pre- vs post-treatment (indicated) with pre-treatment alone compared to post-treatment alone and 24 hr pre-treatment, were compared. J4FUT2-KI HIEs were treated as indicated with 30 uM of TAK-779. HIEs were then infected with GII.3 HuNoV (2.9 × 10^5^ GEs/well) in the indicated conditions. After washing, the cells were cultured for 48 hrs at 37°C in the presence of 30 uM of TAK-779. Viral GEs at 1 and 48 hpi were quantified by RT-qPCR. Mean data compiled from two independent experiments with three wells per experiment are shown; error bars show SD. Experiments are denoted with different symbol colors. Significance was determined using two-way ANOVA comparing 48 h replication, no TAK-779 to each concentration of TAK-779 (P value -**, < 0.001; *“*, 0.0001).

### TAK-779 Enhances GII.3 HuNoV Replication in a Dose- and Time-Dependent Manner and Facilitates Viral Spread

Building on the observed enhancement in GII.3 replication by treating the HIE cultures pre- and post-infection, we next examined whether this effect is time dependent. Pretreatment of HIEs with two concentrations of TAK-779 (15 µM and 30 µM) and assessing viral replication at 24, 48, and 72 hpi showed a significant increase in viral replication with both TAK-779 concentrations compared to the no inhibitor control, with replication increasing in a dose-dependent manner (Fig. 4A). A sharp rise in genome equivalents occurred between 24 and 48 hpi, followed by a plateau at 72 hpi, suggesting that peak replication is achieved by 48 hpi. To determine whether TAK-779 also facilitates viral spread, we performed immunofluorescence analysis on the infected monolayers detecting the viral capsid protein VP1 to quantify HuNoV-infected cells at all time points. The number of infected cells increased with TAK-779 treatment in a dose-dependent manner, further supporting its role in promoting viral spreading (Fig. 4B and 4C). These findings demonstrate that TAK-779 enhances GII.3 HuNoV replication in a dose- and time-dependent manner and facilitates viral spread, representing the most robust replication observed in our system.

**Figure 4.**
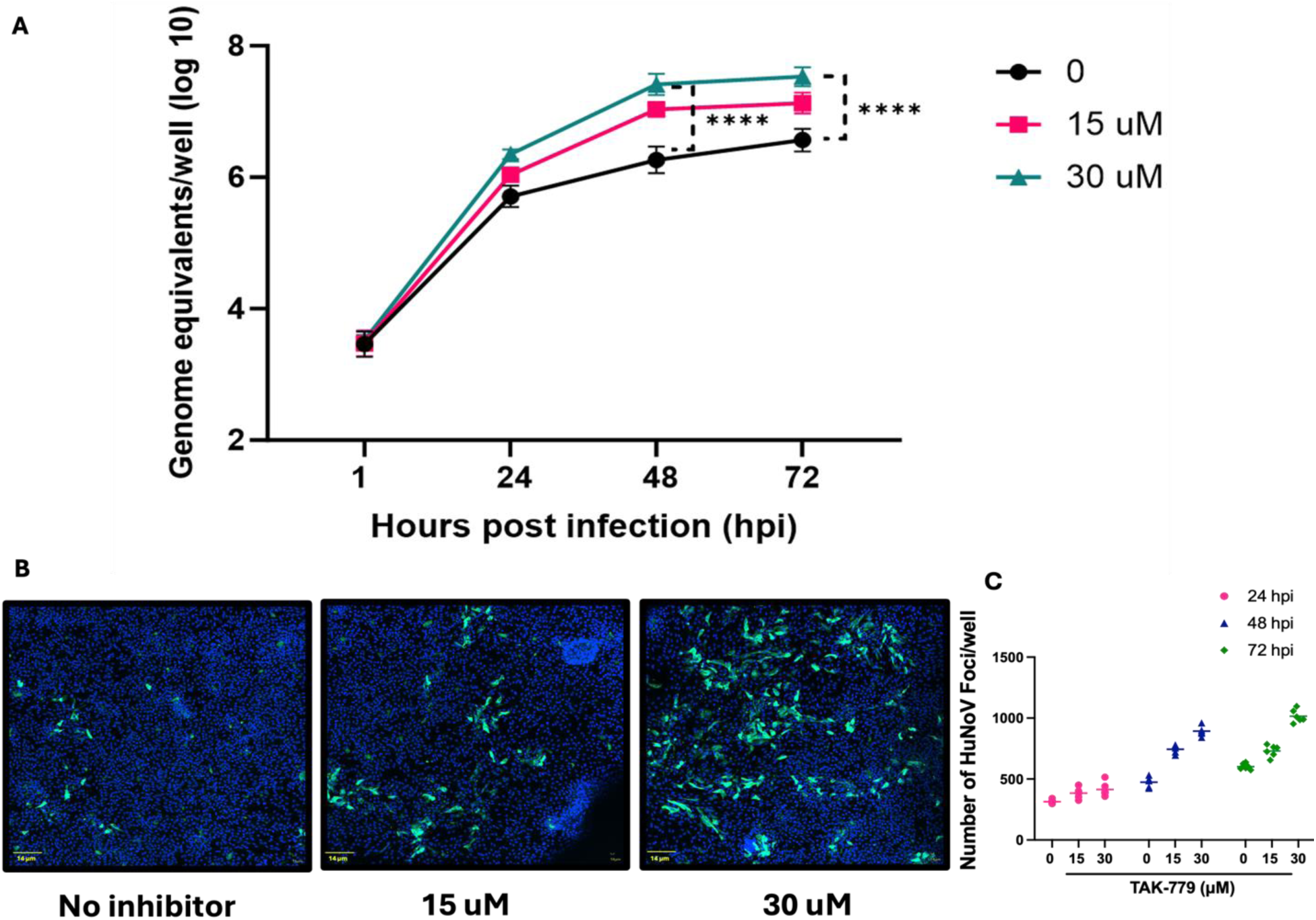
TAK-779 enhances GII.3 replication in a dose- and time-dependent manner and allows viral spreading. (A) J4FUT2-KI HIEs were pre-treated for 3 hrs and then infected with Gil.3 HuNoV (2.9 × 10^5^ GEs/well) in the presence of the indicated concentrations of TAK-779. After washing, the cells were cultured for indicated times at 37°C in the presence of the indicated concentrations of TAK-779. Virus replication was evaluated by RT-qPCR at the indicated time points. (B and C) After 24, 48, and 72 hrs incubation, monolayers were fixed with methanol. Gll.3-positive cells (green) were detected using guinea pig anti-HuNoV VLP Ab (green) and nuclei (blue) were detected by DAPI. (B) Representative images from each condition at 48 hpi are shown. Scale bars denote 14 µm. (C) Numbers of HuNoV infected cells (Foci)/well from each time point and condition were counted. Mean data compiled from two independent experiments with three wells per experiment are shown; error bars show SD. Experiments are denoted with different symbol colors. Significance was determined using one-way ANOVA comparing 48h and 72h replication, 0 to each concentration of TAK-779 (P value -****, 0.0001).

### TAK-779 Treatment Significantly Enhances Viral Infectivity by Reducing the GE/TCID₅₀

To assess whether TAK-779 influences the susceptibility of host cells to GII.3 HuNoV infection, we determined the 50% tissue culture infectious dose (TCID₅₀) in the presence and absence of the chemokine antagonist. The genome equivalents per TCID₅₀ (GE/TCID₅₀) were significantly lower (p <0.05) in TAK-779-treated HIEs compared to the untreated control, indicating that fewer viral genomes were required to establish infection in 50% of the host cells (Table 2). This suggests that, in addition to enhancing viral replication and spread, TAK-779 increases the susceptibility of cells to viral infection. These findings further support the role of chemokine signaling as a critical barrier to HuNoV infection in HIEs, with TAK-779 effectively modulating host restriction factors to favor viral replication and entry.

**Table 2.** GE/TCID_50_ values of GII.3 in the presence and absence of TAK-779. Values represent geometric mean GE/TCID_50_ from n = 3 experiments and expressed as log_10_(GE) required to achieve an infectivity of 50% of the inoculated cultures at 48 hpi. Italicized values in parentheses correspond to standard deviation (SD). Significance was determined using T-test, P-value <0.05.

| HIE – J4FUT2-KI | Log<sub>10</sub> GE/ TCID<sub>50</sub> |  |
| --- | --- | --- |
| HIE culture | + TAK-779 | - TAK-779 |
| Geometric mean (Standard deviation) | 2.99 (0.32) | 3.69 (0.24) |

### TAK-779 Facilitates Sustained Passaging of GII.3 HuNoV in HIEs

Having established that TAK-779 enhances GII.3 HuNoV replication in a dose- and time-dependent manner and promotes viral spread, we next evaluated whether TAK-779 could facilitate sustained viral passaging in two HIE lines, J4FUT2-KI and J8FUT2-KI. In the first round of experiments, we initiated passage 1 (P1) using stool filtrate in the presence of TAK-779 and allowed the infection to progress for up to 96 hours, providing multiple rounds of viral replication. The resulting supernatants were collected, processed as described in the Materials and Methods, and subsequently used for successive passages. Passaging of GII.3 in the absence and presence of TAK-779 was performed in parallel to confirm whether TAK-779 alone is responsible for the ability to passage GII.3 HuNoV. In the absence of TAK-779, viral replication decreased with passaging and ceased at P4 in J4FUT2-KI and P3 in J8FUT2-KI, respectively, whereas in the presence of TAK-779, viral propagation continued up to P5 with sustained virus yields between passages (Fig. 5).

**Figure 5.**
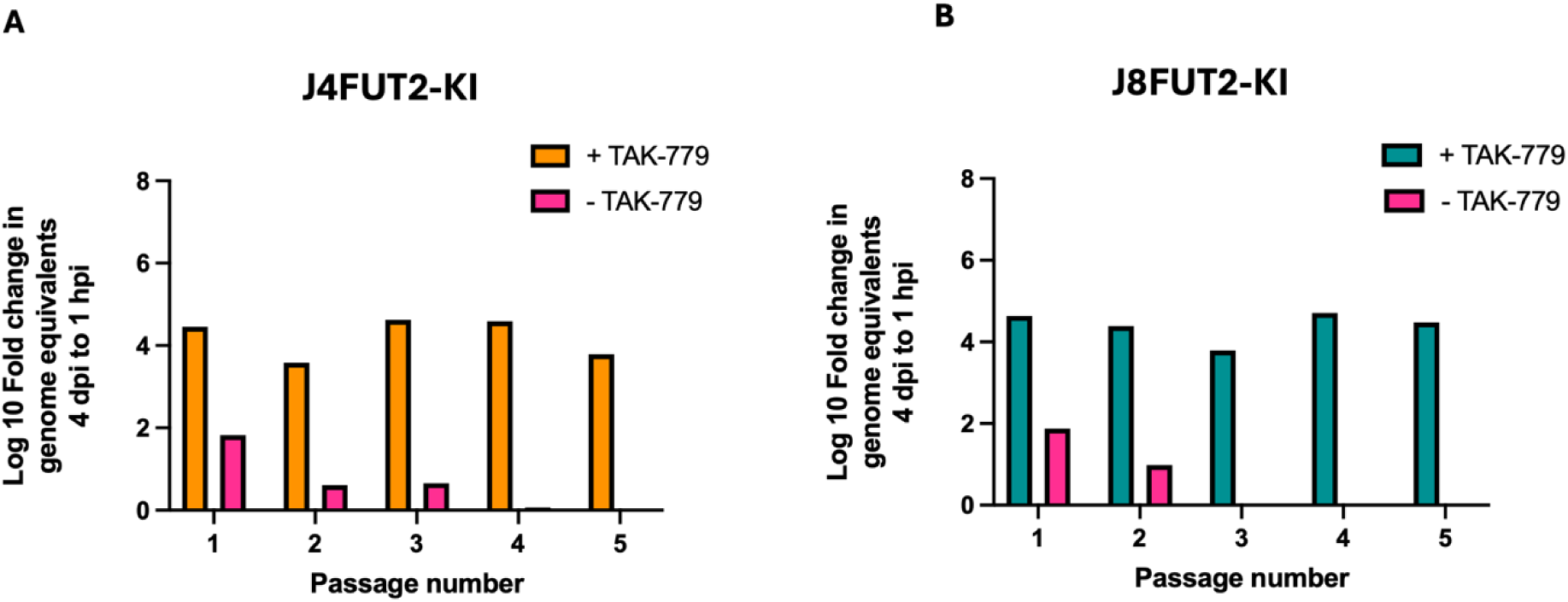
TAK-779 facilitates the passage of GII.3 HuNoV in J4FUT2-KI and J8FUT2-KI cells. (A) J4FUT2-KI and (B) J8FUT2-KI HIEs were pre-treated for 3 hrs and then infected with GII.3 HuNoV (2.9 × 10^5^ GEs/well) with or without 30 uM TAK-779, labeled as P1. Following infection, both cells and supernatant were collected at 96 hpi, and a portion was used to quantify viral RNA by RT-PCR. The virus was subsequently used to infect a fresh batch of HIEs for the next passage, continuing for up to five passages

Building upon these findings, we performed an independent round of passaging in both J4FUT2-KI and J8FUT2-KI cell lines. Remarkably, for the first time, we were able to sustain HuNoV passaging for up to 10 consecutive rounds in both HIE lines, a significant advancement beyond our previous studies (Fig. 6A) (*20*). In J8FUT2-KI, we continued to passage GII.3 up to passage 15 (Fig. S4). To confirm the presence of infectious virus over multiple passages, we performed immunostaining for the VP1 structural and the VPg and NTPase nonstructural proteins at various passages (P2 and P5), demonstrating continued expression of viral structural and nonstructural proteins following infection with the passaged virus (Fig. 6B). These findings highlight the critical role of TAK-779 in allowing the extended HuNoV replication in HIEs, enabling for the first time robust passaging of any HuNoV.

**Figure 6.**
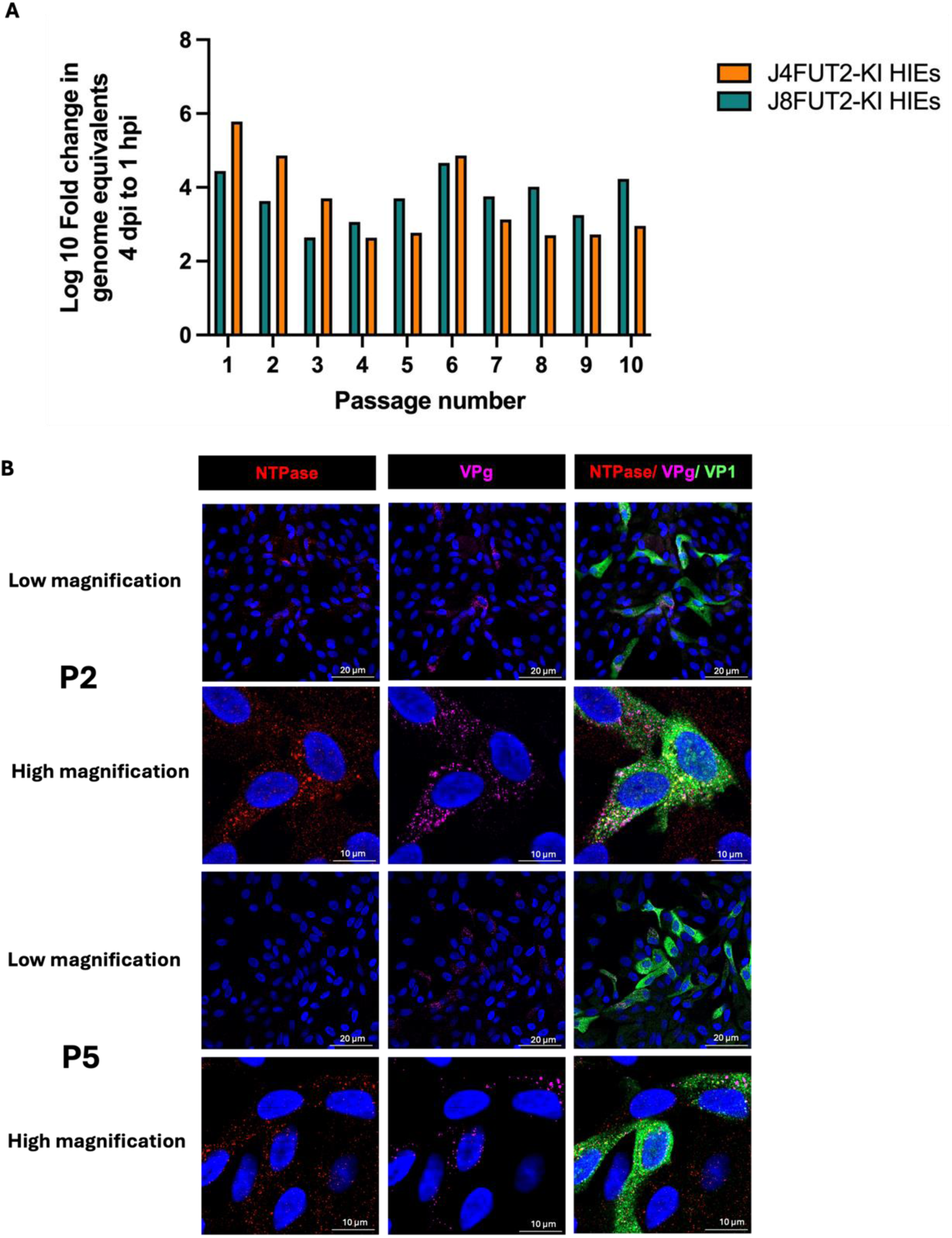
First-time passaging of a HuNoV in HIEs up to 10 passages. (A) J4FUT2-KI and J8FUT2-KI HIEs were pre-treated and then infected with GI1.3 HuNoV (2.9x 105 GEs/well) in the presence of 30 uM TAK-779 and indicated as P1. Post-infection, the cells and supernatant were harvested at 96 hpi; after processing the supernatant, an aliquot was used to quantify the viral RNA by RT-qPCR. The virus was then used to infect a new batch of HIEs for the next passage, continuing up to 10 passages. (B) Immunofiuorescence staining of HuNoVinfected J4AFUT2-KI monolayers at P2 and P5. At 48 hpi, cells were fixed with methanol and stained with guinea pig anti-HuNoV VLP antibody (green), rabbit anti-NTPase (red), and mouse anti-VPg (purple). Nuclei were counterstained with DAPI (blue). Representative images are shown at high (10 ym scale bar) and low (20 um scale bar) magnification.

### GII.3 HuNoV Passaging Enabled Generation of High Titer GII.3 HuNoV Stocks

Following the successful passaging of GII.3 HuNoV in HIEs, we evaluated whether virus collected from different passages could be used as a consistent and scalable source of infectious stocks. J4FUT2-KI HIEs were pre-treated with 30 μM TAK-779 and infected with the original stool-derived GII.3 HuNoV or four independent virus stocks from two different passages. Stock 1 and Stock 2 were derived from P14 virus propagated in J8FUT2-KI HIEs, yielding a total of 15 mL of virus-containing supernatant. Stocks 3 and 4 were generated using P4 virus in J4FUT2-KI HIEs, producing 20 mL total volume. We compared the infectivity of the stocks to GII.3 HuNoV stool infection as an internal control in the presence and absence of TAK-779. In the presence of TAK-779, GII.3 Stocks 1, 2, and 3 exhibited robust replication comparable to the stool-derived virus with similar kinetics (Figure 7A, left panel) indicating replication competence. Stock 4 exhibited lower overall replication compared to the other three stocks but still showed significant infectivity. In the absence of TAK-779, a reduction in replication, approximately 1 log10, was observed across all stocks at 24 hrs, but replication was comparable to stool at later time points 48 and 72 hrs (Figure 7A, right panel). To further validate viral replication, we performed immunofluorescence staining of infected J4FUT2-KI HIE monolayers at 24 hours post-infection. In the presence of TAK-779, all three tested stocks demonstrated widespread infection as indicated by VP1-positive staining (green) distributed across the cell monolayers (Figure 7B). Together, these findings provide a critical advancement for the field by enabling stool-independent propagation of infectious HuNoV material for downstream applications.

**Figure 7.**
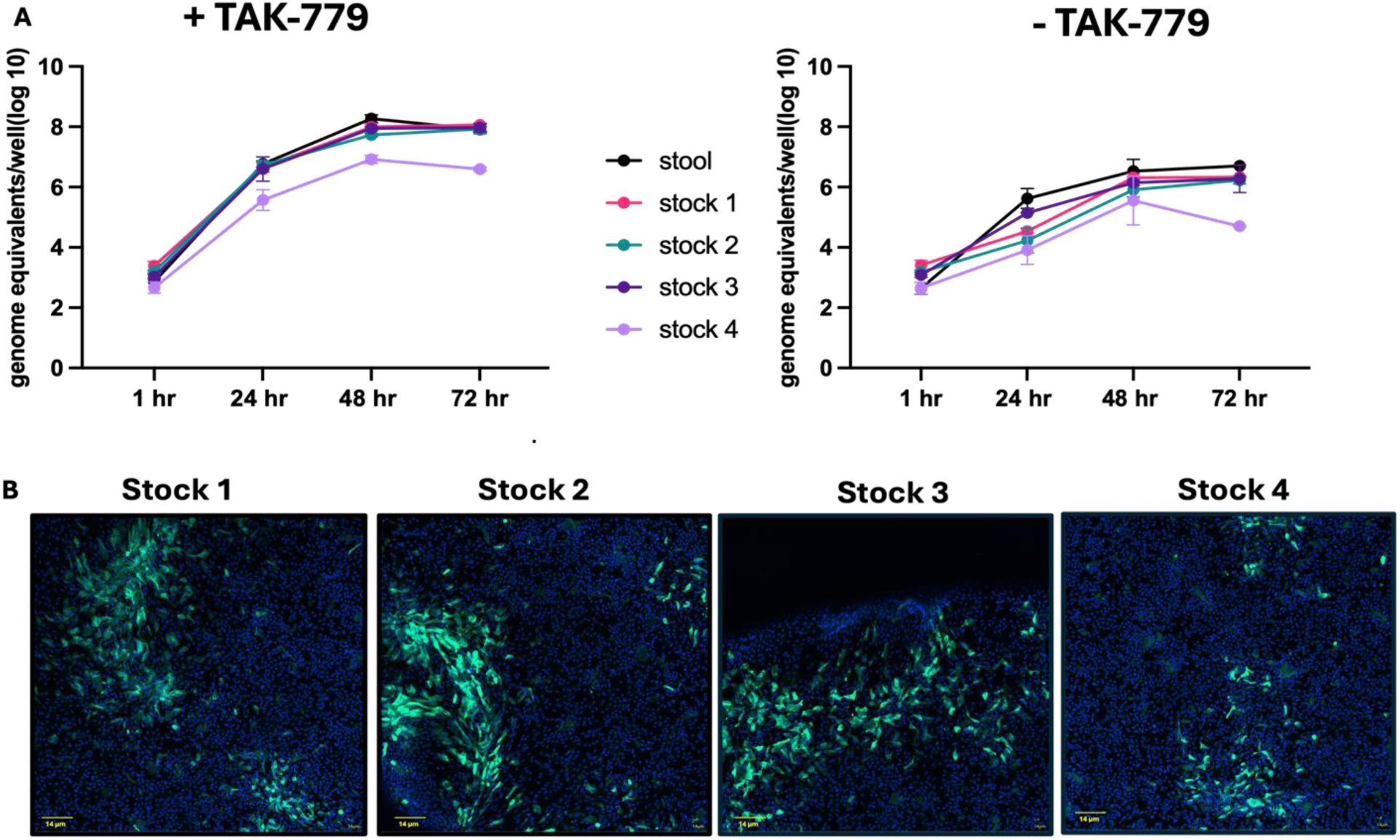
GII.3 HuNoV passaging and TAK-779 enabled generation of Gil.3 HuNoV stocks. (A) J4FUT2-KI HIEs were pre-treated for 3 hours with 30 uM TAK-779 and infected with GII.3 HuNoV derived from stool (2.9 × 10^5^ genome equivalents [GEs]/well) or from Stock 1 (2.58 ×10^5^ GEs/well), Stock 2 (2.93 ×10^5^ GEs/well), Stock 3 (1.02 ×10^5^ GEs/well), or Stock 4 (2.50 × 10^5^ GEs/well). Infections were performed in the presence of 30 µM TAK-779 (left panel) or vehicle control (right panel). After washing, the cells were cultured for the indicated times at 37°C in the presence (left panel) or absence (right panel) of TAK-779. Virus replication was evaluated by RT-qPCR at the indicated time points. Viral replication was assessed by RT-qPCR. The characterization of each stock was performed once with three replicates. Error bars denote standard deviation. (B) Representative images of Stock-infected HIEs fixed with methanol at 48 hpi. GII.3-positive cells were detected using guinea pig anti-HuNoV VLP Ab (green) and nuclei (blue) were detected by DAPI. Scale bars denote 14 µm.

### TAK-779 Enhances GII.17 and GI.1, but not Pandemic GII.4, HuNoV Replication

To further investigate the impact of TAK-779 on HuNoV replication, we evaluated the effect of 30 μM TAK-779, previously identified as the optimal dose, on additional HuNoV genotypes, an emerging GII.17 strain, the GI.1 prototype strain, and a pandemic GII.4 strain, using GII.3 as an internal control. TAK-779 significantly enhanced replication of GII.3 and both GII.17 and GI.1, with greater than 0.5 log₁₀ increases in genome equivalents per well at 48 hpi compared to untreated controls. In contrast, replication of the pandemic GII.4 HuNoV strain remained unaffected by TAK-779 treatment, indicating a lack of response to TAK-779 in this genotype (Fig. 8). This unexpected finding adds a new phenotype to the list of strain-specific differences of HuNoVs (*24, 25, 28, 29, 63*) and indicates that the pandemic GII.4 HuNoV induces a different immune response in HIEs or has mechanisms to antagonize innate immune response.

**Figure 8.**
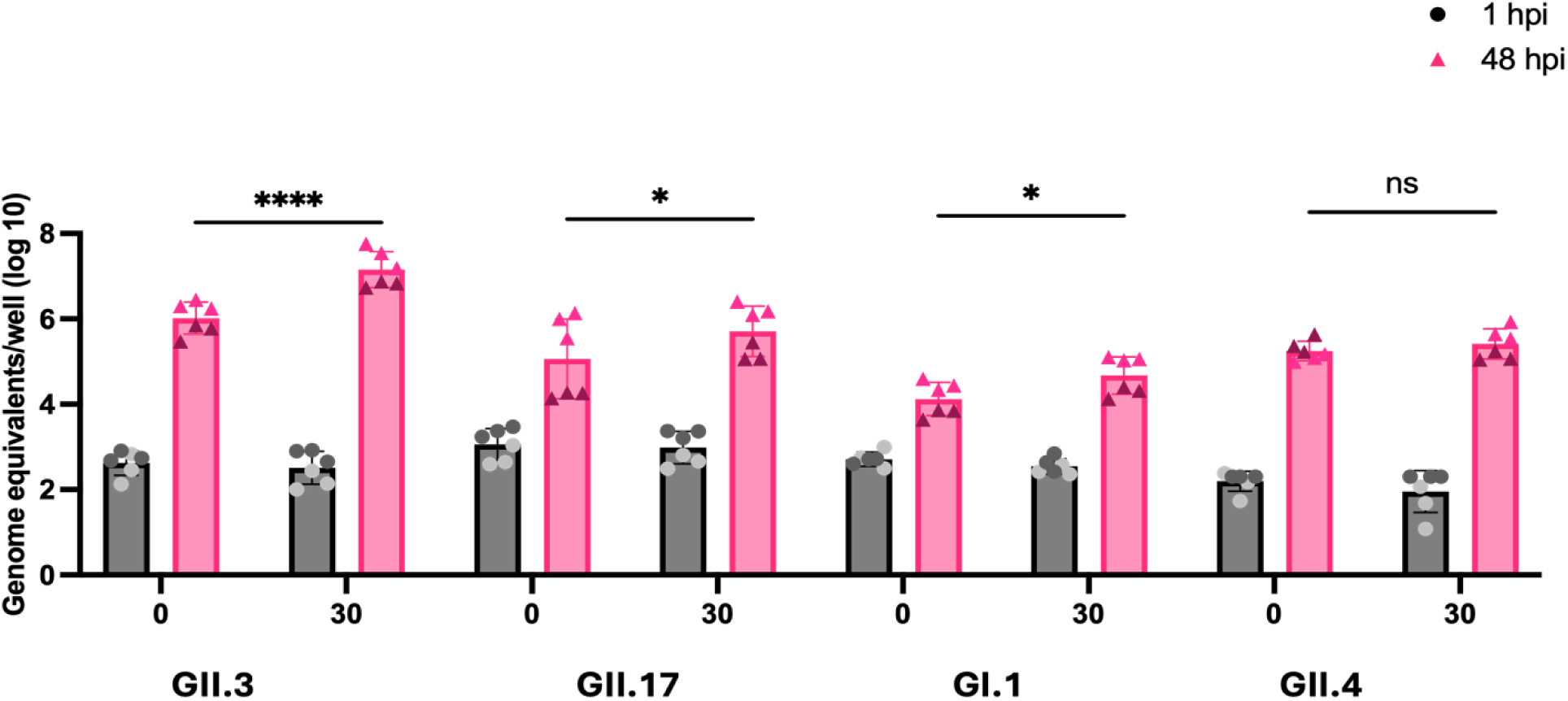
TAK-779 enhances replication of other HuNoV strains GII.17 and G1.1, but not GII.4. J4FUT2-KI HIEs were pre-treated with 30 µM TAK-779 and then infected with GII.3 (2.9 × 10® GE/well), GII.17 (2.8 × 10^5^ GEs/well), G1.1 (3.7× 10^5^ GEs/well), or GII.4 (3.0χ 10^ε^ GE/well) in the absence or presence of 30 µM TAK-779. After washing, the cells were cultured for 48 h at 37°C in the absence or presence of 30 µM TAK-779. Virus replication at 1 and 48 hpi was evaluated by RT-qPCR. Mean data compiled from two independent experiments with three wells per experiment are shown; error bars show SD. Experiments are denoted with different symbol colors Error bars denote standard deviation. Significance was determined using one-way ANOVA comparing 48 h replication, 0 to 30 uM of TAK-779 (P value-*, 0.05; “**, 0.0001).

To further explore the basis of this differential response, we measured the levels of three chemokines, CXCL10, CXCL11, and CCL5, in GII.3- and GII.4-infected cultures using Luminex. While infection with GII.3 infection induced detectable levels of these chemokines, GII.4 infection resulted in no measurable secretion of chemokines (Fig. S5). Poly(I:C) treatment, added as a positive control, induced strong chemokine responses, while mock and gamma-irradiated virus controls showed no induction of chemokines. These results suggest that the inability of TAK-779 to enhance GII.4 replication may be attributed to GII.4-mediated suppression of chemokine synthesis or secretion, highlighting a potential immune evasion strategy unique to this strain.

## Discussion

HuNoVs remain a leading cause of acute gastroenteritis worldwide, yet progress in understanding their biology has long been hindered by the lack of a reliable and scalable in vitro cultivation system. In this study, we overcame a critical barrier in the field by demonstrating that using the small-molecule chemokine receptor antagonist TAK-779 enables robust replication and serial passaging of GII.3 HuNoV in HIEs. By leveraging genetically-modified HIE lines with altered innate immune signaling, we identify a critical role for epithelial chemokines in restricting viral replication. This work establishes a scalable platform for generating high-titer HuNoV stocks, potentially removing a reliance on patient-derived stool samples and reveals a distinct and new component of host antiviral defense beyond interferon signaling.

Historically, HuNoV research has relied on surrogate viruses due to the inability to cultivate the virus efficiently (*64–69*). Recent breakthroughs, however, have led to the development of *de novo* infection models that support the complete viral life cycle and provide new opportunities to study HuNoV replication and biology as well as virus-host interactions in physiologically relevant systems. Various model systems have been explored as cultivation systems for HuNoV. These include B cell lines, tissue stem cell-derived human intestinal enteroids (HIEs), induced pluripotent stem cell (iPSC)-derived intestinal organoids, SV40-transformed human salivary gland cells, and in vivo models such as chimpanzees, gnotobiotic piglets, and zebrafish larvae. Among these, HIEs have emerged as a particularly physiologically-relevant and reproducible human cell cultures platform for studying HuNoV biology. However, the ability to passage HuNoV in vitro remained limited, with most systems supporting only a few rounds of replication. Although GII.17 strains have been reported to be passaged up to eight times in iPSC-derived HIOs, that study did not address further passaging or the potential to generate high-titer virus stocks (*18*). Other systems, zebrafish larvae and salivary gland cell lines, have supported replication up to four passages but exhibited decreased replication with each passage, similar to our previous observations in the absence of TAK-779. As a result, the field has lacked an ex vivo culture system capable of generating high-titer virus stocks and studies have remained dependent on patient-derived stool samples (*16, 17, 19*)

Herein, we probed our genetically-modified J4FUT2-KI and J2STAT1-KO HIE lines to begin to understand why they support enhanced replication of multiple virus strains (*22, 29,*). RNA-seq analysis revealed that both modified lines exhibited a dampened interferon response to GII.3 infection. While J2 cells mounted a strong interferon-stimulated gene (ISG) response, the modified lines showed reduced ISG induction but maintained high expression of epithelial chemokines, including CXCL10, CXCL11, and CCL5. Notably, J4FUT2-KI, engineered to express FUT2 in a non-secretor background (*22, 24*) also exhibited attenuated interferon signaling with a concomitant strong upregulation of chemokines including CXCL10, CXCL11, CCL20, CXCL1, and CXCL2. Together, these findings highlighted the potential role of epithelial chemokine signaling as a distinct antiviral barrier in HuNoV infection.

The upregulation of CXCL10 has been documented in multiple previous HuNoV infected HIE studies (*27, 29, 71, 72*) as well as in two persons naturally infected with HuNoV (*73, 74*). In early controlled human infection studies with the prototype GI.1 HuNoV, elevated serum CXCL10 was detected early and remaining elevated after infection (*75, 76*). In another clinical study, an early (2 days after HuNoV infection) increase of CXCL10 and other chemokines was detected in a single patient (*73*). A clinical study of hospitalized patients with GII.4 HuNoV gastroenteritis reported elevated serum levels of CXCL10, CXCL9, IL-18, soluble IL-2 receptor, and macrophage inhibitory factor (MIF), with low CCL5 levels correlating with prolonged viral shedding—suggesting a role for CCL5 in viral clearance (*74*). In our study, GII.3 HuNoV infection induced significant upregulation of both CCL5 transcripts and protein levels in HIEs, supporting its potential role in antiviral immunity. Additional data from multiple studies in norovirus-infected HIEs that profiled innate responses confirmed the importance of interferon and chemokine upregulation, often focusing on both gene and protein expression of CXCL10 (*27, 29, 71, 72*). Together, these data underscore the contribution of chemokine-mediated signaling to host defense mechanisms during HuNoV infection.

Given the strong chemokine response in modified HIEs, we tested TAK-779, primarily known to target CCR5 and block the replication of R5-HIV strains in immune cells but it is also known to inhibit CXCR3 and CCR2 (*62, 77–81*). We found unexpected and remarkable enhancement in GII.3 HuNoV replication in three HIE lines, J2, J4FUT2-KI and J8FU2T-KI, with TAK-779 treatment. Increased viral spreading and culture susceptibility to infection (based on lower TCID50) in the presence of TAK-779, was accompanied by continued, successful passaging of GII.3 HuNoV for 10-15 rounds of passaging in HIEs. This success has translated into the successful generation of GII.3 HuNoV stocks that can now be used for detailed mechanistic, structural, and biochemical studies. Our findings have the potential to reduce the reliance on stool samples for inocula and enable laboratories lacking access to clinical isolates to perform HuNoV research. Importantly, the extended passaging was only achieved in the presence of TAK-779, highlighting the critical role of chemokine signaling in limiting sustained replication.

These results raise several new questions that we are exploring. First, it is important to determine if passaging the virus results in selection of virus that contains specific mutations or adaptive changes that may have occurred during serial culture. Since, TAK-779 also enhances GII.17 and GI.1 HuNoV replication, it will be important to evaluate if these and other strains are able to be passaged in the presence of this compound. By contrast, we observed no replication enhancement with the pandemic GII.4 strain, which notably failed to induce detectable protein levels of chemokines such as CXCL10, CXCL11, and CCL5. These findings suggest that a possible explanation for the inability of TAK-779 to enhance GII.4 replication may stem from the capacity of GII.4 HuNoV to suppress the synthesis or secretion of host chemokine or degrade chemokines, underscoring a potential strain-specific mechanism of immune evasion. Understanding how GII.4 circumvents these responses will provide valuable insight into strain-specific immune evasion strategies.

Our findings open the door to obtaining high-purity stocks for structural studies of live, infectious GII.3 HuNoV. This would provide information on whether there are structural differences between virus-like particles (VLPs) and virions containing viral RNA and inform the rational design of antiviral agents. The exact mechanism by which TAK-779 enhances replication in HIEs also remains to be determined. Identification of chemokine receptor(s) expressed on specific cells in HIE cultures and the precise chemokines being blocked by TAK-779 need to be characterized. Identifying the specific chemokine receptors involved should enable the development of targeted receptor knockout HIE lines that are more permissive to infection. Future experiments using unbiased transcriptomics and proteomics will help uncover the effect of TAK-779 on cells and virus infection. While chemokines are well characterized in immune cells, their role in virus-infected epithelial cells is less understood. Our immune co-culture models will allow us to further dissect the interplay between chemokines, epithelial cells, and the immune system during HuNoV infection.

## Acknowledgements

This research was supported by National Institutes of Health Grant P01 AI57788, U19 AI116497, and P30 DK56338 that supports the Texas Medical Center Digestive Diseases Core Center, S10 OD028480 that supported purchasing the Zeiss Laser Scanning Microscope LSM 980 with Airyscan 2.

