## Supplement for "Overcoming host restrictions to enable continuous passaging of human noroviruses in human intestinal enteroids"

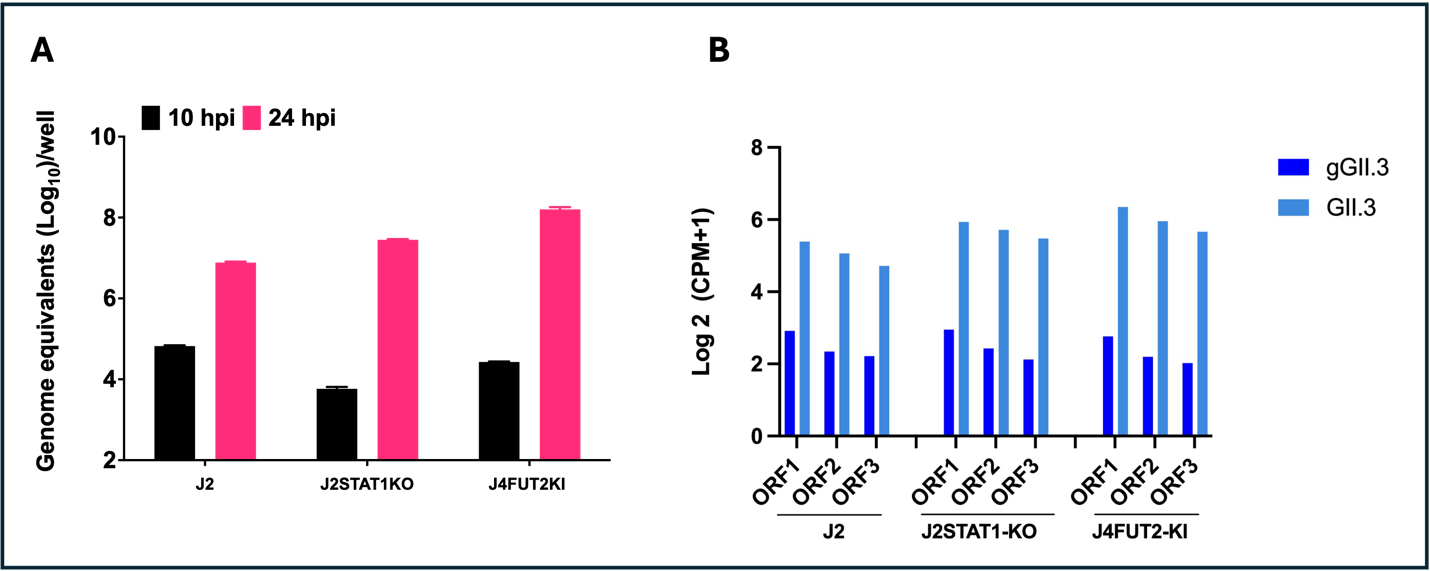

**Supplemental Figures**

Supplemental Figure 1. GII.3 HuNoV replication in HIEs and detection of GII.3 ORFs1-3 in RNA-seq samples from different HIEs. (A) Differentiated HIE monolayers were inoculated with GII.3 (3.2 x10^7^ GE/well) in presence of 500 μM GCDCA. After 1 hpi, monolayers were washed twice and cultured in Intesticult differentiation medium (+ 500 μM GCDCA). Black bars indicate GE at 10 hpi and pink bars show GE at 24 hpi. Error bars show SD. (B) RNA Sequencing data was mapped using STAR to a reference comprised of the human genome and the GII.3 genome sequence, then gene expression was quantified using feature counts. Gene expression, normalized to counts per million (CPM), is shown for ORF1, ORF2 and ORF3, in log2(CPM+1) scale. Gene abundance is shown for gamma-irradiated virus inoculated- (dark blue bars) and in GII.3-infected HIEs (light blue bars).

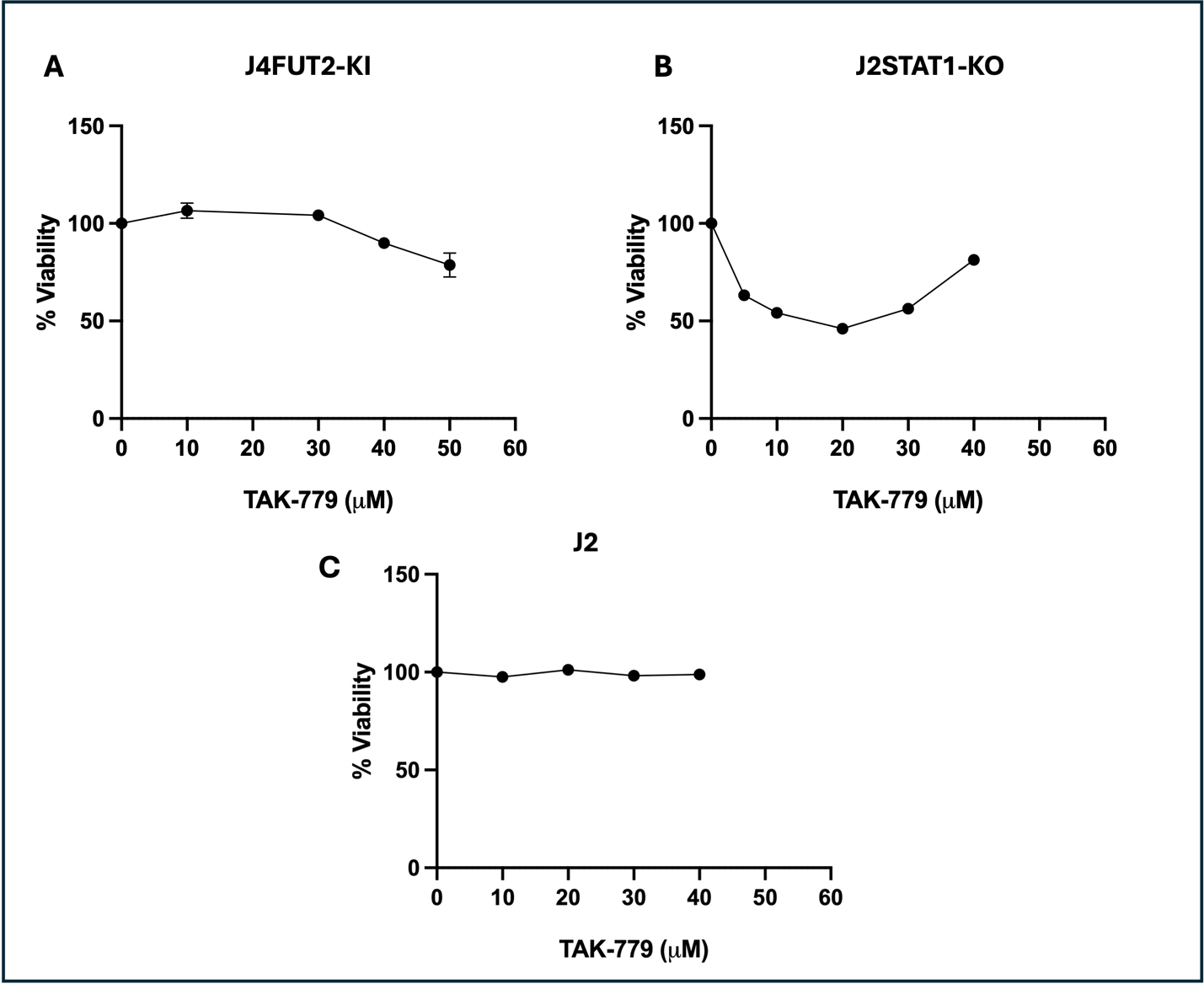

Supplemental Figure 2. TAK-779 exhibits cytotoxicity above 30 μM in J4FUT2-KI and at lower concentrations in J2STAT1-KO HIEs with no cytotoxicity in J2 (A) J4FUT2-KI and (B) J2STAT1-KO (C) J2 (HIEs) were treated with varying concentrations of TAK-779 for 48 hrs. Untreated cells were lysed with 0.8% Triton X-100 as a positive control for maximum lysis. Supernatants were collected, diluted 2-fold, and subjected to an LDH release assay using the CytoTox 96® Kit. Cytotoxicity was quantified and plotted using non-linear regression. Data is representative from one experiment performed with two biological wells. Error bars show SEM (Standard error of mean)

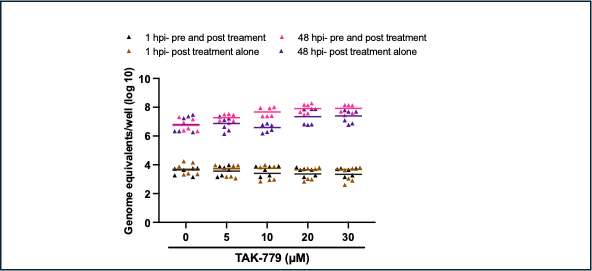

Supplemental Figure 3. Pre- and post-treatment with the CXCR5/CXCR3/CCR2 antagonist TAK-779 results in enhanced GII.3 HuNoV replication compared to pre- or post-treatment alone.
J4FUT2-KI HIEs were treated as indicated with increasing concentrations of TAK-779 for 3 hours or left untreated. HIEs were then infected with GII.3 HuNoV (2.9 × 10⁵ GEs/well) in the presence (pre-treated cells) or absence (untreated cells) of TAK-779. Following infection, monolayers were washed and cultured for 48 hours at 37°C as indicated with TAK-779. Viral replication was assessed by quantifying GEs at 1 and 48 hpi using RT-qPCR. Data are representative from one of two experiments performed with three technical replicates.

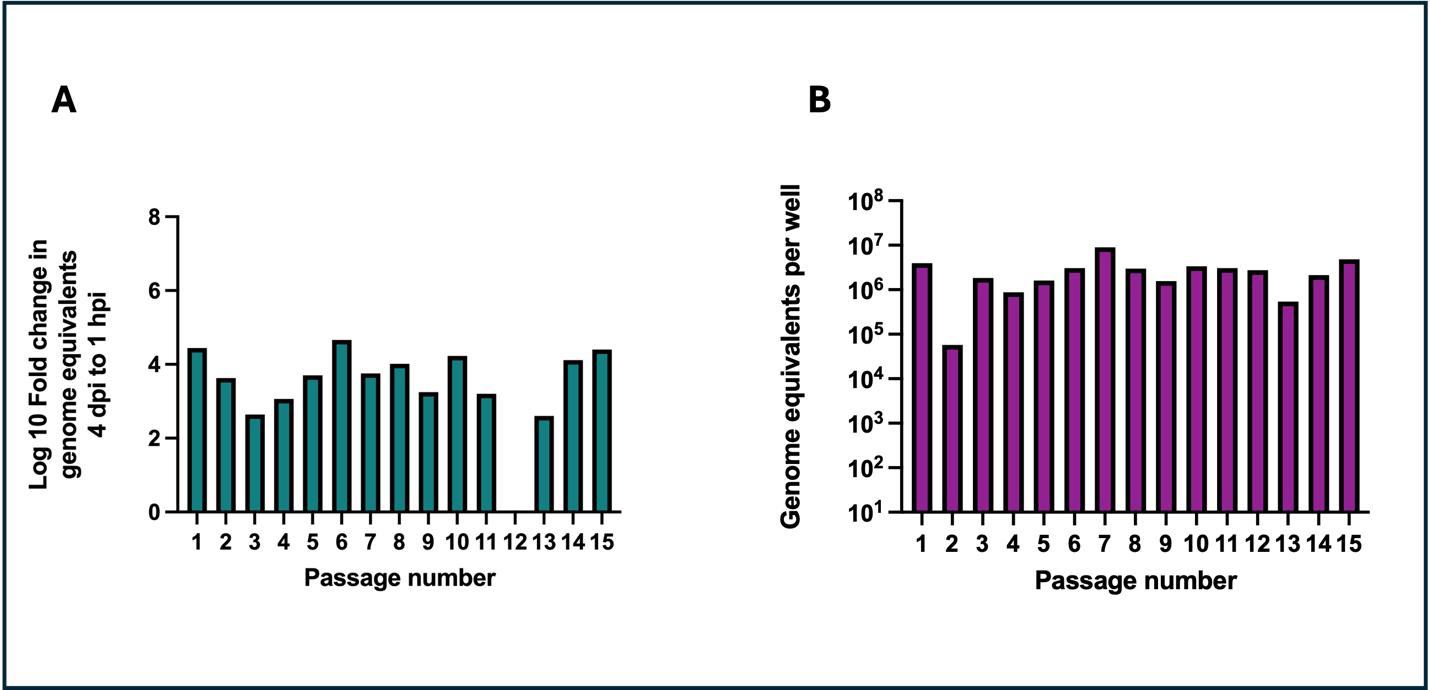

Supplemental Figure 4. Serial passaging of GII.3 HuNoV in J8FUT2-KI HIEs for up to 15 passages in the presence of TAK-779. (A) Log₁₀ fold change in genome equivalents between 1 hpi and 96 hpi for each passage; data for P12 was not collected. (B) Heat release measured at 96 hpi across all passages. J8FUT2-KI human intestinal enteroids (HIEs) were pretreated with 30 µM TAK-779 and infected with GII.3 HuNoV (2.9 × 10⁵ genome equivalents [GEs]/well) indicated as P1. At 96 hours post-infection (hpi), cells and supernatants were harvested, processed, and viral RNA was quantified by RT-qPCR. Supernatants from each passage were used to infect fresh monolayers, continuing the process for a total of 15 sequential passages in the continued presence of TAK-779.

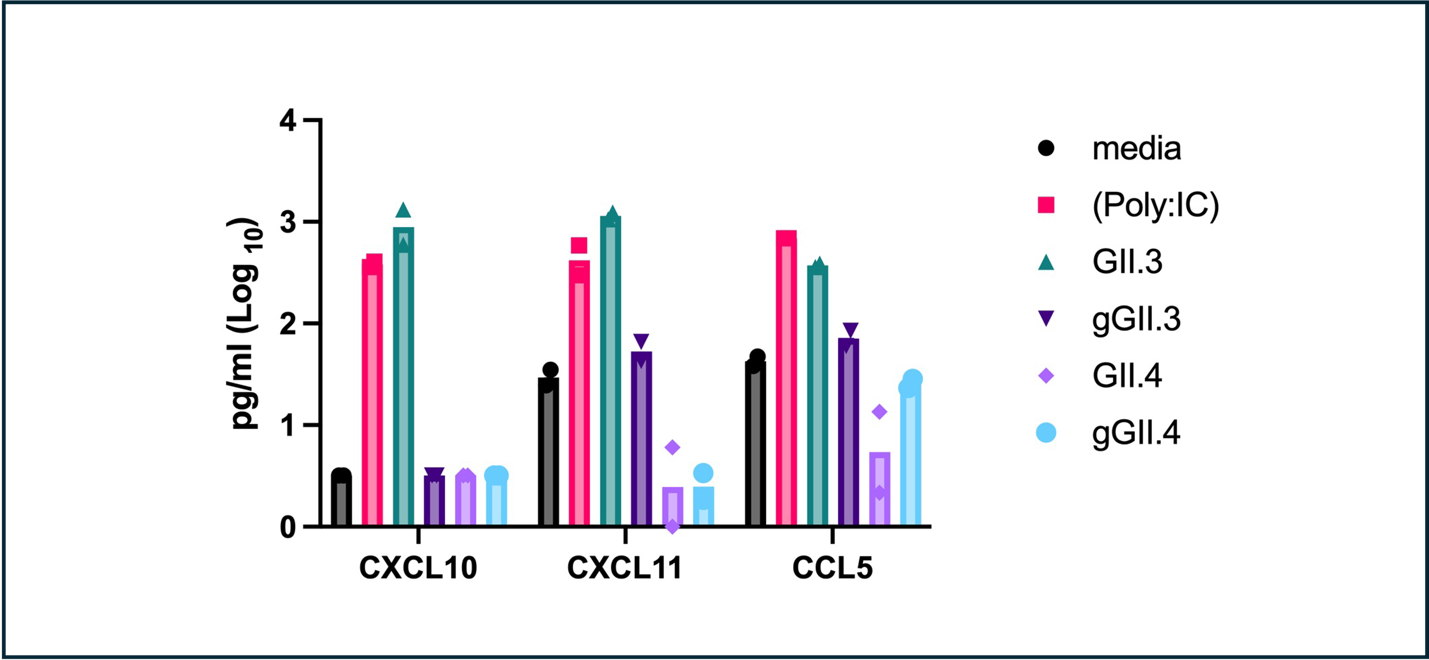

Supplemental Figure 5. Chemokine secretion is induced by GII.3 HuNoV infection but not by pandemic GII.4 HuNoV infection. J2 HIEs were inoculated with media control, GII.4 HuNoV (9 × 10⁷ GEs/well), GII.3 HuNoV (1.7 × 10⁷ GEs/well), poly(I:C) (100 μg/mL), or gamma-irradiated GII.4 or GII.3 HuNoV in the presence of 500 μM GCDCA. Supernatants were collected at 72 hours post-infection (hpi) and analyzed for CXCL10, CXCL11, and CCL5 protein levels [pg/mL (Log _10_)] by Luminex assay. Data represent results from two wells per condition.
